# The Effect of Transcutaneous Auricular Vagus Nerve Stimulation (taVNS) on Cardiorespiratory Physiology: Four Waveforms Trial with Tragus vs Earlobe Stimulation

**DOI:** 10.64898/2026.07.29.741582

**Authors:** Devin Adair, Jacob Itzhakov, Henry Berstein, Mohamad Fallah Rad, Helen Borges, Marom Bikson

**Affiliations:** Department of Biomedical Engineering, City College of New York, New York, NY, USA

## Abstract

**Background:** Transcutaneous auricular vagus nerve stimulation (taVNS) depends on activating afferent vagal pathways, leading to central and physiological (parasympathetic) modulation. Explaining how taVNS changes cardiorespiratory physiology requires systematic characterization of the effects of electrode location and stimulation waveform using rigorous experimental controls.

**Objective:** To determine the acute effects of four taVNS stimulation configurations (left unilateral 25 Hz, 100 Hz, burst, and bilateral 25 Hz) on physiological markers of parasympathetic activation compared with configuration-matched earlobe stimulation in healthy subjects at rest.

**Methods:** Customized 8-mm ear-clip electrodes delivered monophasic 500 μs pulses (anode anterior) to the tragus or a configuration-matched earlobe control across four stimulation configurations (left unilateral 25 Hz, 100 Hz, burst, and bilateral 25 Hz). Twenty-five healthy participants completed a randomized, single-blind, within-subject crossover study, undergoing both active and control stimulation for all four configurations (200 sessions total). Each session consisted of five 60-second stimulation blocks (1,000 stimuli total). Heart rate (HR), heart rate variability (HRV; RMSSD), and respiration were recorded continuously. Linear mixed-effects models compared physiological responses between tragus and earlobe stimulation during the first stimulation block (primary analysis), across 5-second HR intervals (secondary analysis), and averaged across all five stimulation blocks (post hoc analysis).

**Results:** During the first stimulation block, HRV was significantly higher during unilateral left 25 Hz tragus stimulation than during the matched earlobe control (β = −10.34, t(33) = −2.63, p = .01, 95% CI [−18.05, −2.64]), consistent with increased parasympathetic activity. No significant differences were observed for HRV with the other stimulation configurations, or for HR or respiration under any configuration. Analysis of 5-second HR intervals likewise revealed no significant temporal effects for block one. Analysis of 5-second heart rate intervals across the five blocks identified differences between tragus and earlobe at discrete time points but did not show a consistent temporal pattern across the stimulation period. Across all five stimulation blocks, unilateral left 25 Hz tragus stimulation remained associated with higher HRV (β = −6.56, t(124) = −3.85, p < .01, 95% CI [−9.90, −3.22]) and also produced an increase in HR relative to earlobe stimulation (β = −1.05, t(24) = −2.57, p = .01, 95% CI [−1.85, −0.24]). Respiration was unaffected throughout.

**Conclusion:** In this systematic study, unilateral left 25 Hz tragus stimulation produced physiological modulation consistent with vagal target engagement. These findings underscore the importance of rigorous dose-specific taVNS studies to establish reproducible physiological biomarkers.

## Introduction

Transcutaneous auricular vagus nerve stimulation (taVNS) is noninvasive electrical stimulation of the external ear, intended to activate afferent fibers of the auricular branch of the vagus nerve, thereby producing central and physiological modulation. Afferent vagal fibers project to brainstem regions, including the nucleus tractus solitarius (NTS), which in turn modulates parasympathetic autonomic activity (Chen and Liu, 2025). Consequently, heart rate (HR), heart rate variability (HRV), and respiration are typical biomarkers of taVNS target engagement. Despite expanding clinical interest, the physiological mechanisms of taVNS and the stimulation parameters that reliably activate vagal afferents remain incompletely understood (Wienke et al., 2023)

A mechanistic understanding of taVNS requires systematic characterization of the stimulation parameter space. Stimulation dose is defined not simply by current amplitude but also by electrode location, frequency, and waveform that together determine neural recruitment (Badran et al., 2019; Beynel et al., 2019; Dias et al., 2024). Yet few studies have directly compared these parameters (Atanackov et al., 2025; Badran et al., 2019; Farmer et al., 2021; Kreisberg et al., 2021; Sclocco et al., 2020a). Instead, ad hoc stimulation protocols and control conditions have complicated comparisons across studies and limited understanding of which parameter combinations most effectively engage vagal pathways (Wienke et al., 2023).

To examine key dimensions of taVNS dose, we evaluated four active stimulation configurations that varied waveform frequency, temporal patterning, and laterality. The conventional unilateral left-ear 25 Hz configuration represents the most common taVNS paradigm in both clinical and experimental studies (Farmer et al., 2021; George et al., 2000). Bilateral 25 Hz stimulation tested whether increasing the stimulated auricular territory enhanced autonomic responses, the 100 Hz configuration evaluated frequency-dependent recruitment, and a burst configuration examined whether temporal patterning influenced physiological responses. For each configuration, tragus stimulation was compared with a waveform-matched earlobe control to isolate vagal-specific effects from nonspecific somatosensory stimulation.

The objective of this study was to determine how stimulation location and configuration influence the physiological effects of taVNS in healthy adults at rest. Using a randomized, subject-blind, within-subject crossover design, we compared four tragus stimulation configurations with waveform-matched earlobe controls while continuously recording heart rate (HR), heart rate variability (HRV), and respiration. Because autonomic responses to taVNS may evolve with repeated stimulation, physiological outcomes were evaluated during both the initial stimulation block and across all five stimulation blocks. The initial block was used to characterize the immediate physiological response to stimulation onset, whereas analyses across all five blocks assessed whether these responses were maintained, diminished, or strengthened with repeated stimulation. Heart rate was additionally analyzed in 5-second intervals to determine whether time-averaged analyses masked short-lived physiological responses during stimulation. We hypothesized that tragus stimulation would elicit greater parasympathetic activation than matched earlobe stimulation, reflected by decreased HR, increased HRV, and decreased respiration, and that these responses would depend on the stimulation configuration.

## Methods

The experiment was designed to investigate the effects of different taVNS dosages on physiological measures. A description of the experimental method follows:

### Participants

Twenty-nine healthy adults (9 women, 20 men) were enrolled following approval from the City College of New York Institutional Review Board, and all participants provided written informed consent. Participants were excluded if they had a history of cardiovascular disease, seizure disorders, head or ear trauma, were taking medications known to affect cardiac function, or were unable to tolerate the stimulation device. Completion of all eight experimental sessions was required for inclusion in the final analyses. Four participants did not complete all sessions and were excluded, leaving a final sample of 25 participants.

### Experimental Design and Procedure

The study employed a randomized, subject-blind, within-subject crossover design to examine the physiological effects of taVNS. Each participant completed eight experimental sessions, four active tragus stimulation configurations and four configuration-matched earlobe control conditions. Sessions were separated by at least 24 hours.

Prior to testing, participants completed questionnaires assessing demographics, previous electrical stimulation experience, sleep quality, and adherence to a required 2-hour abstinence from food and caffeine. Participants were seated in a sound-attenuated room illuminated only by the computer display with their head stabilized in a chin rest. ECG electrodes and a respiratory effort belt were placed and calibrated, the stimulation site was disinfected with alcohol, and stimulation intensity was determined using the threshold procedure described above.

Each session began with a 150-second baseline recording while participants viewed a fixation cross (Figure 1). Five identical stimulation blocks followed, each consisting of a 15- second visual countdown, 60 seconds of stimulation, 60 seconds of post-stimulation recording, an on-screen pain rating, and a 30-second rest period. Sessions were terminated if pain ratings reached ≥5; however, no participant met this criterion.

**Figure 1.**
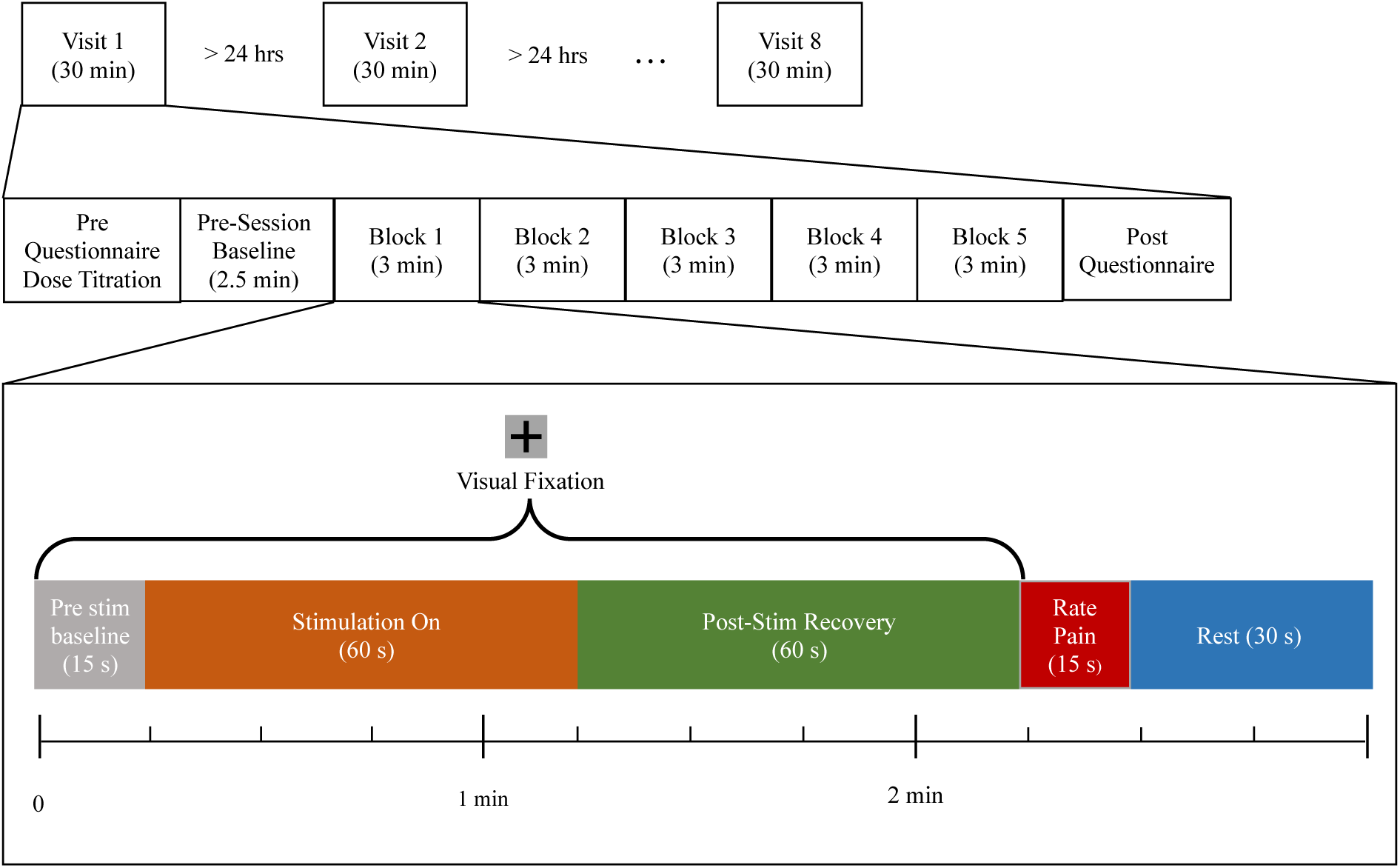
Schematic overview of the experimental protocol and block design. The session began with a 150-s baseline period during which participants fixated on a central cross while stabilizing their head in a chin rest. Each stimulation block started with a 15-s countdown, followed by 60 s of electrical stimulation and a 60-s post-stimulation interval. Participants then rated stimulation pain (0–10) using a number pad, after which a 30-s rest period was provided. Five stimulation blocks were completed per visit. After the final block, participants completed post-experiment questionnaires (≈15 min), yielding a total session duration of ∼30 min.

The stimulation protocol required approximately 30 minutes, with total visit duration of approximately one hour including instrumentation, questionnaires, and adverse event reporting.

### Stimulation Protocol

Four stimulation configurations were evaluated (Figure 2a): unilateral left 25 Hz, unilateral left 100 Hz, burst stimulation (five 100 Hz pulses repeated at 5 Hz), and bilateral 25 Hz stimulation. Pulse width was held constant at 500 μs for every configuration and stimulation location.

**Figure 2.**
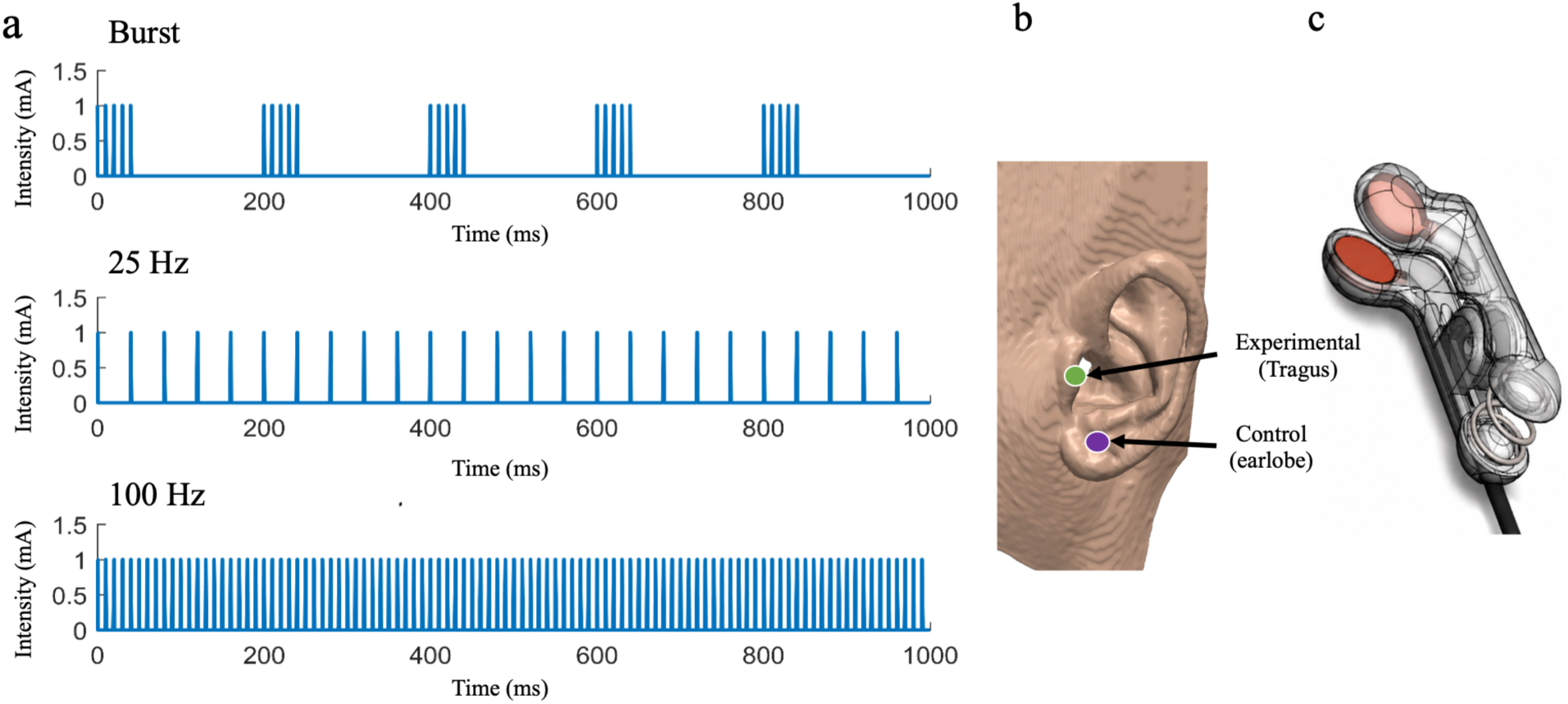
Electrical stimulation waveforms, location and device. a) Waveforms: Each pulse has a width of 500μs. The burst waveform grouped five 100 Hz pulses at 5 Hz. The 100 Hz and 25 Hz waveforms delivered pulses at 100 Hz and 25 Hz, respectively. The unshown 25 Hz pulsed bilateral waveform is identical to the 25 Hz waveform but applied to both ears simultaneously. b) Location of the active electrode (tragus, green) and control electrode (earlobe, purple). c) The clip features 8 mm copper-coated bipolar electrodes within a 3D-printed resin body, coated with Ten20 conductive paste.

**Figure 3.**
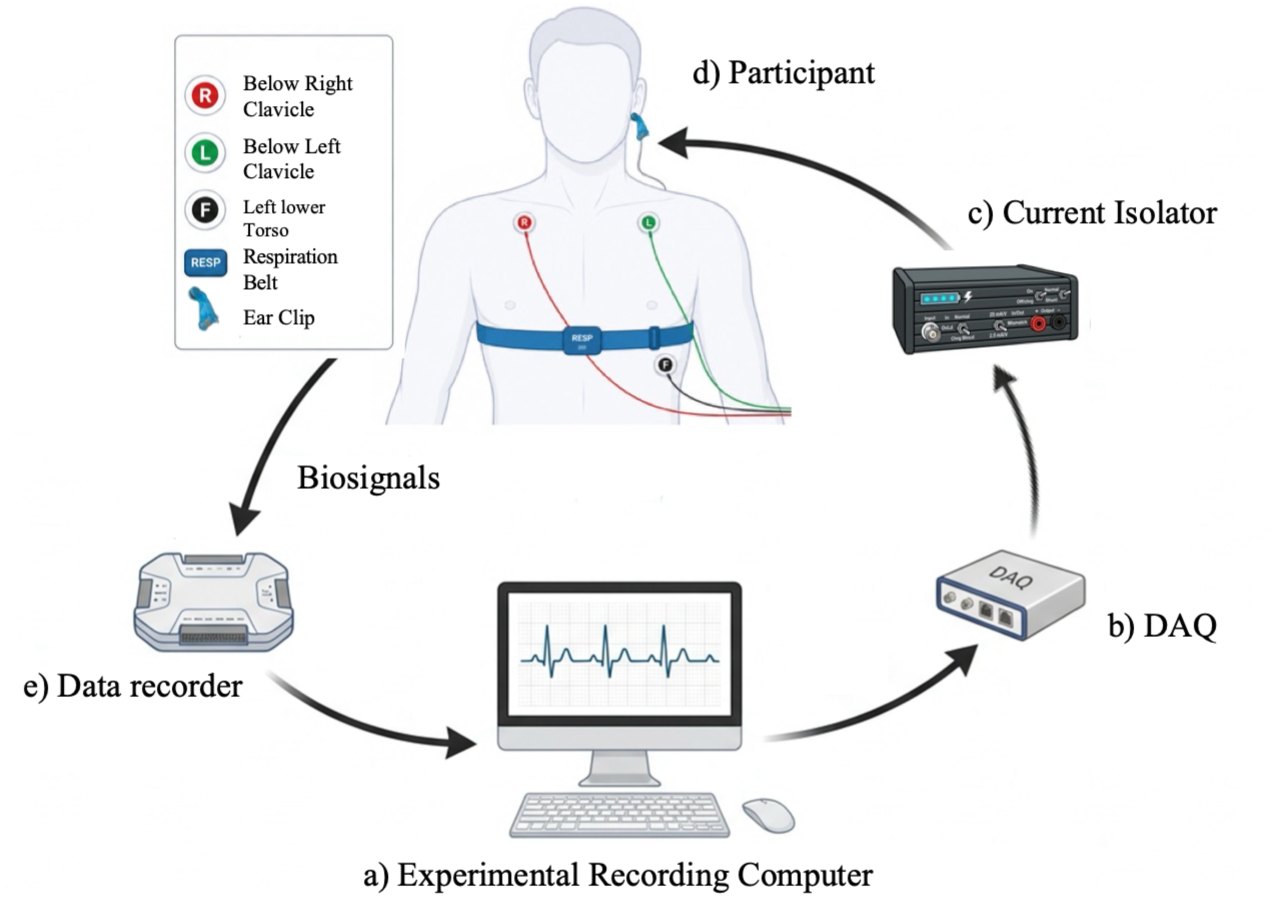
Experimental tech setup for acquiring biosignals data and applying electrical stimulation a) Experimental recording computer b) DAQ accepts digital signals from the experimental computer c) Current isolator receives an analog signal from the DAQ d) The participant receives electrical stimulation on the ear while ECG and respiration are collected e) All physiological signals are transmitted into the data recording device and then transmitted back to the experimental computer.

The unilateral 25 Hz configuration represents the conventional stimulation paradigm most used in taVNS research. The 100 Hz configuration evaluated frequency-dependent effects, the burst configuration examined temporal patterning, and bilateral stimulation tested whether increasing the spatial footprint of stimulation altered physiological responses.

For each configuration, stimulation was delivered either to the tragus (active) or to a configuration-matched earlobe control (Figure 2b).

The tragus was selected because it contains dense innervation from the auricular branch of the vagus nerve (Badran et al., 2018; Peuker and Filler, 2002; Safi et al., 2016). The earlobe served as the active control because it contains minimal vagal innervation while preserving the somatosensory experience of electrical stimulation (Fernandez-Hernando et al., 2023).

Unilateral configurations were delivered to the left ear based on conventional clinical practice. Bilateral stimulation was included to evaluate whether increasing the spatial extent of stimulation altered autonomic responses.

Electrical stimulation was delivered using a custom-designed spring-loaded ear clip fabricated in-house. The clip contained two opposing 8-mm diameter copper contacts (anode anterior) housed within a 3D-printed electrically insulated body constructed from PLA/ABS filament. Electrode cups were filled with conductive paste (Ten20 Conductive Paste; Weaver and Company) immediately before application to maintain stable electrode impedance.

To accommodate differences in ear anatomy, interchangeable compression springs (McMaster-Carr) provided varying clamping forces. Spring selection was individualized during the initial session to provide stable electrode contact while maintaining participant comfort.

Device tolerance was assessed immediately following fitting using a Visual Analog Scale (VAS). A pain score ≥ 5 on the VAS was deemed unacceptable, requiring spring adjustment to ensure the clip could be tolerated comfortably for the duration of the 30-minute experimental protocol.

Electrical stimulation was delivered via an in-house current isolator (Figure). Using our own stimulator design provided a wider range of dose parameters than off-the-shelf equipment and offered lower cost than some commercially available products. The circuit is based on the Improved Howland current pump with bridge output and floating load. The voltage-to-current conversion is implemented using OPA145 op amps and 0.01%, 1/4-W precision resistors. The OPA145 features a low offset of 40 μV, a high open-loop gain of 123 dB, and a gain-bandwidth product of 5.5 MHz. The isolator’s compliance voltage is about ±36 V, with an input range of ±10 V and an output impedance of 30 MΩ. The isolator’s linear accuracy is 0.01%, with two selectable ranges of 100 μA/V and 1 mA/V, corresponding to feedback resistor values of 500 Ω and 5 kΩ, respectively. The isolator can minimize the potential for even small, undesired DC currents (Merrill et al., 2005) during precise neurostimulation experiments. The LCI’s power source consists of 3 Li-Ion cells in series, each with a 3400 mAh capacity, battery protection circuitry, charge circuitry, and a dual-output DC/DC boost converter followed by passive filters and accurate linear LDO regulators.

### Stimulation Intensity and Thresholding

Stimulation intensity was individualized before every session using a perceptual threshold procedure (Cornsweet, 1962). Single 500 μs monophasic pulses were delivered beginning at 2 mA and progressively reduced until no longer perceived. The final perceptual threshold was determined after four staircase repetitions, and experimental stimulation intensity was set to twice this threshold.

Immediately before each session, participants received a 5-second test stimulation at the assigned intensity. If pain was rated ≥5 on a 0–10 numerical rating scale, stimulation intensity was reduced by 25% (up to two times). Participants unable to tolerate stimulation following adjustment were excluded.

Pain ratings were obtained immediately after thresholding, following each stimulation block, and at the conclusion of each experimental session using a 0–10 numerical rating scale. Participants also completed post-session questionnaires assessing physiological symptoms and overall well-being.

### Physiological Recording

Cardiac activity was recorded using a three-lead ECG sampled at 2 kHz with an EEGO-Sport amplifier (ANT Neuro, The Netherlands). Disposable Ag/AgCl electrodes were positioned bilaterally below the clavicles with ground and reference electrodes placed on the torso.

Heart rate (beats/min) and heart rate variability (RMSSD) were derived from the ECG recordings. HRV was calculated over each 60-second stimulation period and baseline-corrected using the pre-session 90-second baseline window. Heart rate was analyzed both across the entire stimulation period and within consecutive 5-second epochs and baseline-corrected for the 15 s before stimulation began for each block. Both Block 1 and all five blocks were analyzed.

Respiration was recorded using a piezoelectric respiratory effort belt connected to the EEGO-Sport amplifier. Signals were band-pass filtered (0.1–0.5 Hz, second-order Butterworth, zero phase), and respiration rate was determined using automated peak detection. Respiration values were calculated over each 60-second stimulation period and baseline-corrected using the pre-session 90-second baseline window

### Statistical Analyses

Perceptual thresholds, stimulation intensity, and pain ratings were compared between tragus and earlobe stimulation within each configuration using paired-samples t-tests with Bonferroni correction (α = 0.0125).

Physiological data were screened for implausible values (HR <30 or >200 bpm, HRV <20 or >100 ms, respiration <12 or >20 breaths/min), which were replaced with missing values prior to analysis. Cardiac and respiratory outcomes were analyzed using linear mixed-effects models. Separate models were fit for each stimulation configuration, with stimulation location (tragus vs. earlobe) specified as the fixed effect and participant as a random intercept. Primary analyses examined responses during the first stimulation block to minimize potential habituation effects. Post hoc analyses evaluated responses averaged across all five stimulation blocks using additional random intercepts for stimulation block.

Heart rate was additionally analyzed across twelve consecutive 5-second epochs using linear mixed-effects models with fixed effects for stimulation location and time and random intercepts for subject and stimulation block. Analyses were conducted for the initial stimulation block and repeated using data averaged across all five stimulation blocks. Pairwise comparisons between stimulation locations at each epoch were performed using estimated marginal means with adjustment for multiple comparisons.

Data preprocessing was performed in MATLAB (R2025a). Statistical analyses were conducted in R using lme4 and lmerTest, with Satterthwaite approximations used to estimate degrees of freedom. Statistical significance was evaluated using two-tailed tests with a Bonferroni-corrected significance threshold of α = 0.0125. Unless otherwise stated, all analyses use tragus as the reference.

## Results

Physiological responses were analyzed according to a planned statistical analysis framework. The primary analysis examined average responses during the 60-second stimulation period of the initial stimulation block (Block 1) to assess the acute effects of taVNS. Because heart rate can fluctuate throughout stimulation, a secondary analysis examined HR in consecutive 5-second epochs to characterize its temporal dynamics. A post hoc analysis evaluated average responses across all five stimulation blocks to assess the effects of repeated stimulation. This analytical framework was applied to heart rate, heart rate variability, and respiration, as appropriate.

### Participants

Four of the 29 enrolled participants withdrew before completing all eight sessions. The final analysis included 25 participants (9 women; age 25.16 ± 4.16 years).

### Perceptual Threshold, Dose, and Tolerability

No adverse events were observed during the experimental sessions or reported following completion of the trial. Transient erythema at the stimulation site resolved within minutes following stimulation.

Perceptual threshold, stimulation intensity (following titration), pre-stimulation pain, and mean pain across the five stimulation blocks were compared between tragus and earlobe stimulation for each configuration (Table 1). No significant differences in perceptual threshold or pain ratings were observed between stimulation sites for any configuration.

**Table 1.**
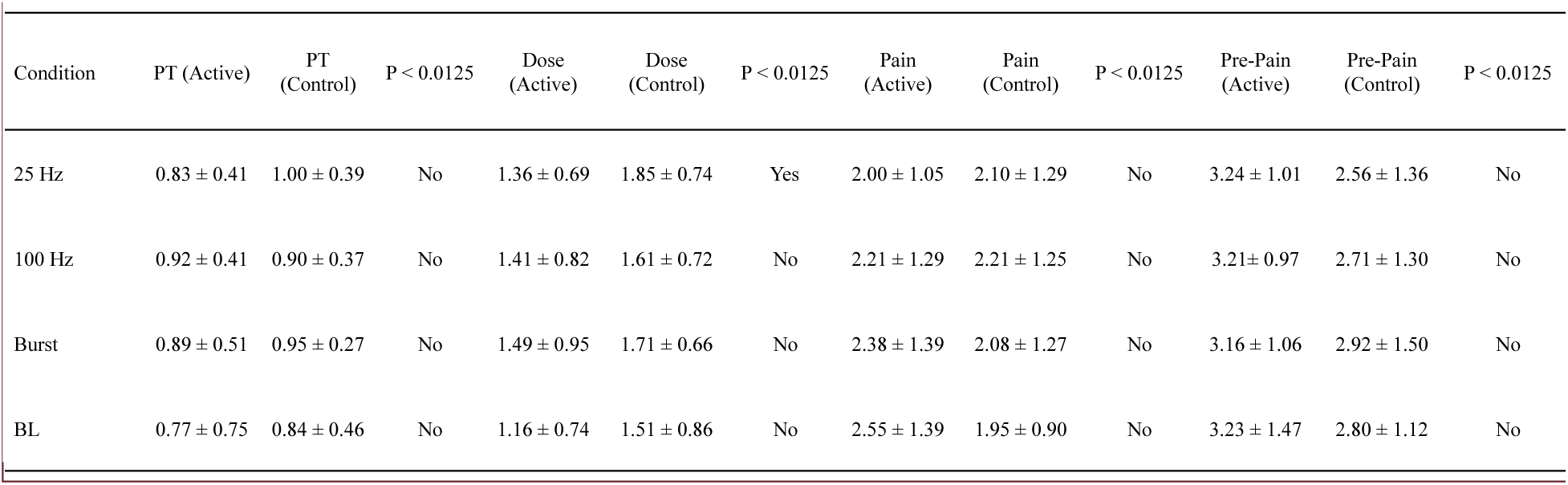
Perceptual threshold, dose and pain levels for each waveform. Each entry is mean (SD). Active is the taVNS condition (electrode on the tragus) and control is the active control condition with the electrode placed on the earlobe. PT is the perceptual threshold (mA). Dose is the stimulation intensity used in the experimental paradigm (mA). Dose starts at 2x the PT and is reduced by 25% (administered at most twice) for any pain rating of 5 or above on a test dose. Pain is rated on an NRS from 0-10. “Pain” is the average pain rating across all five blocks of stimulation. “pre-pain” is the rating given to the 5s test before the start of the experiment.

| Condition | PT (Active) | PT (Control) | P < 0.0125 | Dose (Active) | Dose (Control) | P < 0.0125 | Pain (Active) | Pain (Control) | P < 0.0125 | Pre-Pain (Active) | Pre-Pain (Control) | P < 0.0125 |
| --- | --- | --- | --- | --- | --- | --- | --- | --- | --- | --- | --- | --- |
| 25 Hz | 0.83 ± 0.41 | 1.00 ± 0.39 | No | 1.36 ± 0.69 | 1.85 ± 0.74 | Yes | 2.00 ± 1.05 | 2.10 ± 1.29 | No | 3.24 ± 1.01 | 2.56 ± 1.36 | No |
| 100 Hz | 0.92 ± 0.41 | 0.90 ± 0.37 | No | 1.41 ± 0.82 | 1.61 ± 0.72 | No | 2.21 ± 1.29 | 2.21 ± 1.25 | No | 3.21 ± 0.97 | 2.71 ± 1.30 | No |
| Burst | 0.89 ± 0.51 | 0.95 ± 0.27 | No | 1.49 ± 0.95 | 1.71 ± 0.66 | No | 2.38 ± 1.39 | 2.08 ± 1.27 | No | 3.16 ± 1.06 | 2.92 ± 1.50 | No |
| BL | 0.77 ± 0.75 | 0.84 ± 0.46 | No | 1.16 ± 0.74 | 1.51 ± 0.86 | No | 2.55 ± 1.39 | 1.95 ± 0.90 | No | 3.23 ± 1.47 | 2.80 ± 1.12 | No |

Following dose titration, the unilateral 25 Hz configuration required a significantly lower stimulation intensity at the tragus (1.36 ± 0.69 mA) than at the earlobe (1.85 ± 0.74 mA; t(24) = −3.50, p = .002). No significant dose differences were observed for the remaining configurations.

### Heart Rate: Time Averaged Effects for Block One (Primary Analysis)

HR averaged over the 60-second stimulation period was analyzed separately for each configuration using linear mixed-effects models with stimulation location as a fixed effect and participant as a random intercept (Figure 4). There was no significant effect for any tested stimulation configuration: 25 Hz (β = −1.87, t(24) = −1.87, p = .07, 95% CI = [−3.81, 0.08]), 100 Hz (β = 0.22, t(24) = 0.22, p = .83, 95% CI = −1.69, 2.12]), bilateral 25 Hz (β = −1.04, t(24) = −1.27, p = 0.22, 95% CI = [−2.65, 0.57]), or burst stimulation (β = 0.64, t(48) = 0.79, p = 0.43, 95% CI = [−0.94, 2.23]).

**Figure 4.**
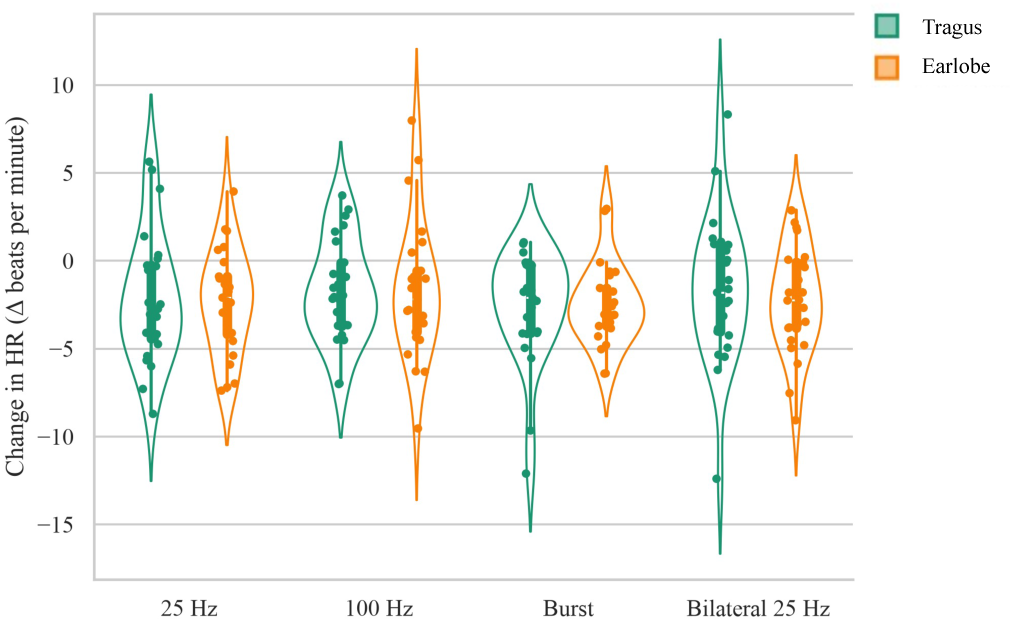
The effects of taVNS at the tragus versus the earlobe on heart rate for Block 1 only. Measured as change from pre-stimulation baseline across four distinct stimulation waveforms: 25 Hz, 100 Hz, Burst, and Bilateral 25 Hz. Data are color-coded by stimulation location: tragus (green) and earlobe (orange). Individual data points represent unique participants, while the width of the violin contours illustrates the probability density of the dataset.

### Heart Rate: Temporal Dynamics for Block One (Secondary Analysis)

To determine whether transient effects were obscured by time averaging, HR was analyzed in twelve consecutive 5-second epochs using linear mixed-effects models with fixed effects for location and time, and a random intercept for subject (Figure 5). No significant differences between tragus and earlobe stimulation were detected at any time point for any stimulation configuration.

**Figure 5.**
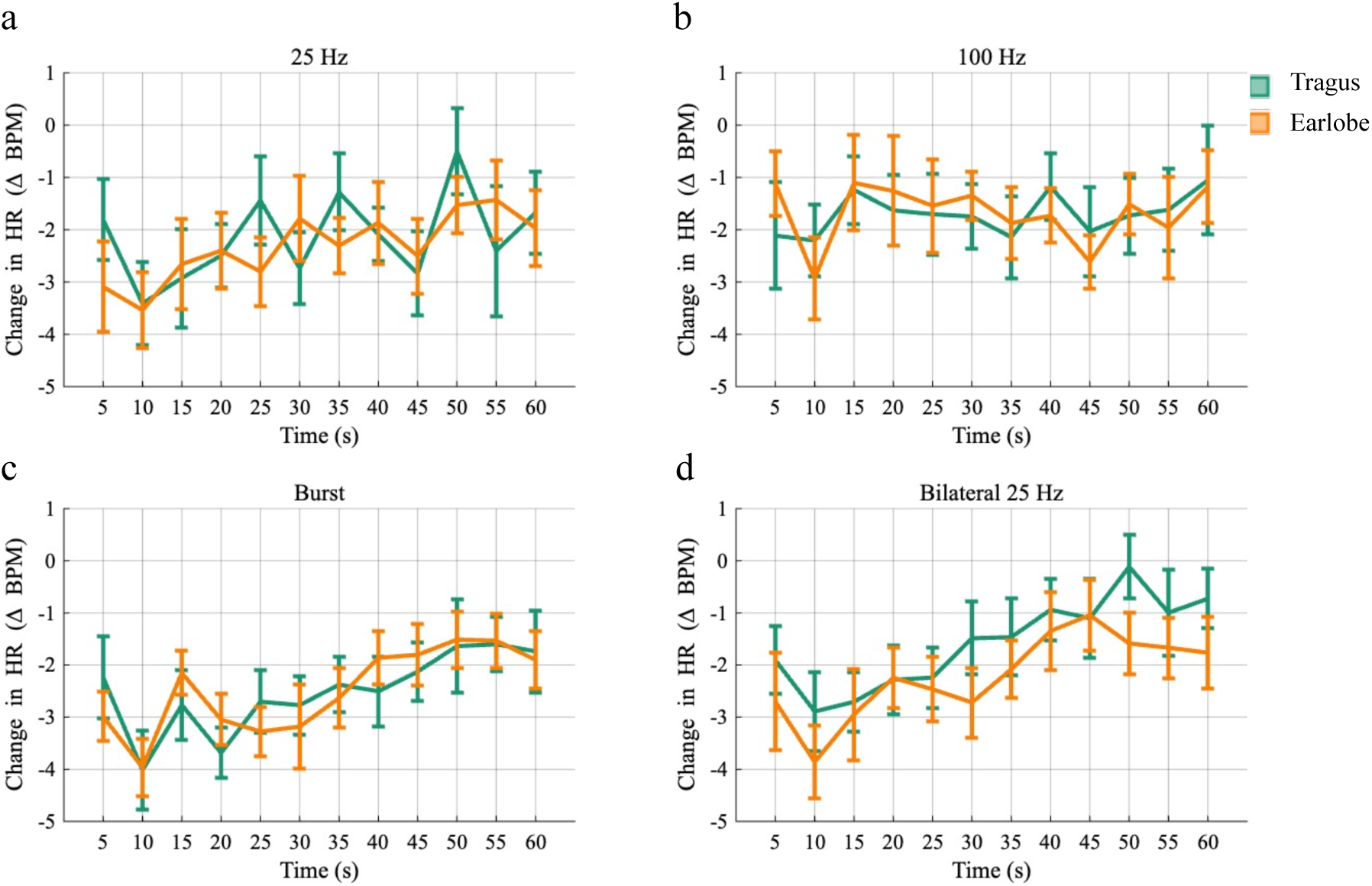
Effects of taVNS waveform on the temporal dynamics of heart rate for block 1. Comparing active stimulation and matched control conditions across 5-second time bins over the 60-s stimulation window. All data are baseline-corrected relative to the pre-stimulation baseline period. a) 25 Hz b) 100 Hz c) Burst d) Bilateral 25 Hz. (Tragus; green) (earlobe; orange).

### Heart Rate: Temporal Dynamics Across Five Blocks (Secondary Analysis)

To evaluate whether physiological responses varied over the stimulation period, HR was analyzed in twelve consecutive 5-second epochs using linear mixed-effects models with fixed effects for stimulation location and time and a random intercept for subject and block within subject (Figure 6). Significant differences between tragus and earlobe stimulation were observed at specific time points for unilateral left 25 Hz (20, 25, 35, 45, 50, and 60 s; Table S1), unilateral left 100 Hz (35 and 60 s; Table S2), and bilateral 25 Hz (40 s; Table S3). No significant differences were observed for the burst configuration (Table S4). This secondary analysis suggests taVNS effects vary during the stimulation period.

**Figure 6.**
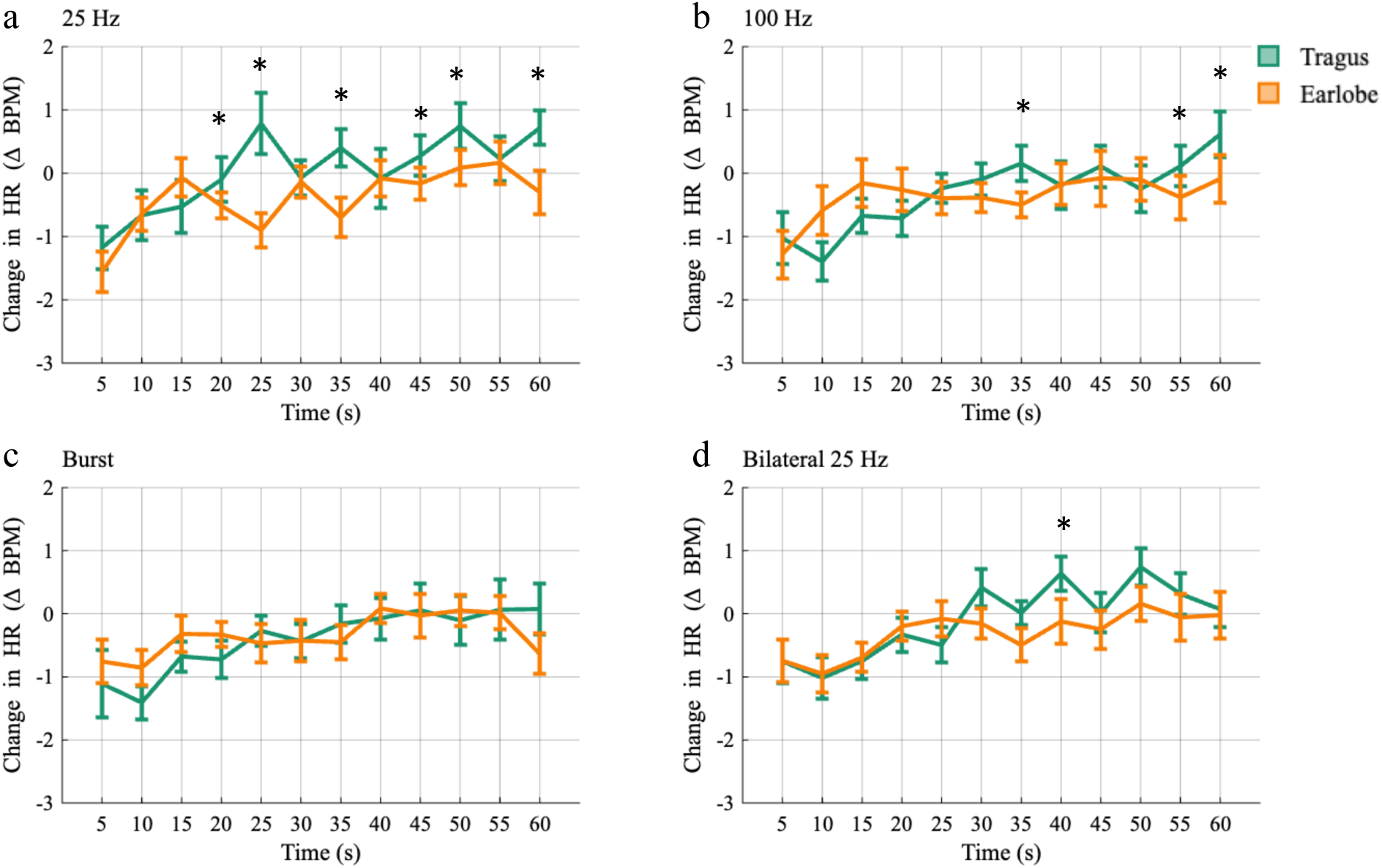
Effects of taVNS waveform on the temporal dynamics of heart rate across five blocks. Comparing active stimulation and matched control conditions across 5-second time bins over the 60-s stimulation window. All data are baseline-corrected relative to the pre-stimulation baseline period. a) 25 Hz b) 100 Hz c) Burst d) Bilateral 25 Hz. (Tragus; green) (earlobe; orange).

### Heart Rate: All Five Blocks (Post Hoc)

HR averaged over the 60-second stimulation period and across all five blocks was analyzed separately for each configuration using linear mixed-effects models with fixed effects for stimulation location and random intercepts for subject and block within subject (Figure 7). A significant effect of stimulation location was observed only for the unilateral 25 Hz configuration (β = −1.05, t(24) = −2.57, p = .01, 95% CI [−1.85, −0.24]). There was no significant effect for 100 Hz (β = −0.70, t(224) = −1.67, p = .10, 95% CI = −1.51, 0.13]), bilateral 25 Hz (β = 0.66, t(124) = −1.82, p = .07, 95% CI = [−1.38, 0.05]), or burst stimulation (β = 0.44, t(224) = 1.05, p = .29, 95% CI = [−0.38, 1.26]).

**Figure 7.**
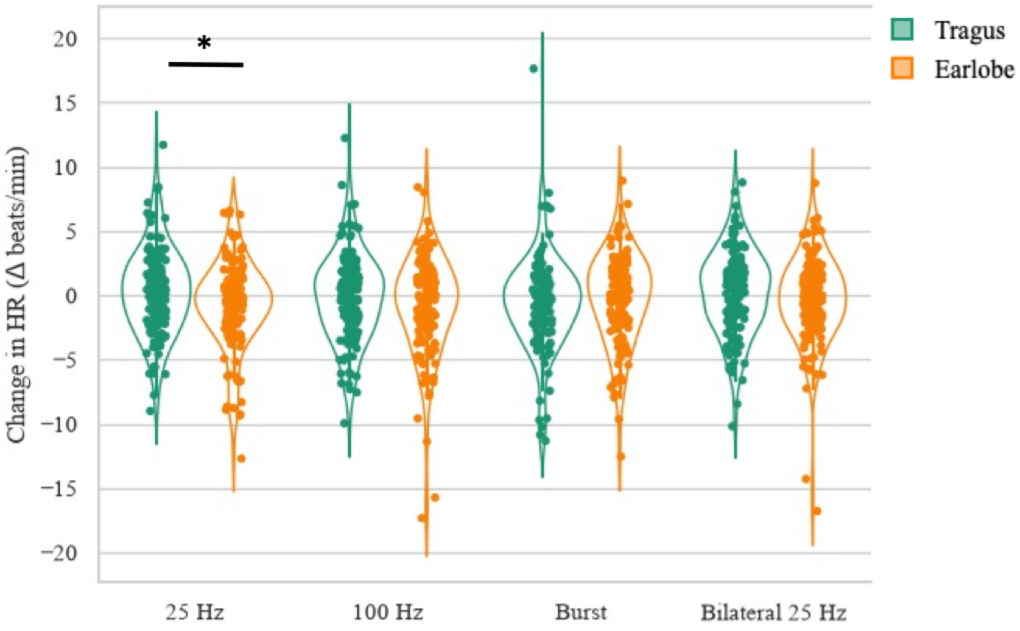
The effects of taVNS at the tragus versus the earlobe on heart rate for all five blocks. Measured as change from pre-stimulation baseline across four distinct stimulation waveforms: 25 Hz, 100 Hz, Burst, and Bilateral 25 Hz. Data are color-coded by stimulation location: tragus (green) and earlobe (orange). Individual data points represent unique participants, while the width of the violin contours illustrates the probability density of the dataset.

### Heart Rate Variability: Time Averaged Effects for Block One (Primary Analysis)

HRV averaged over the 60-second stimulation period was analyzed separately for each configuration using linear mixed-effects models with fixed effects for stimulation location and a random intercept for subject (Figure 8). A significant effect of stimulation location was observed only for the unilateral 25 Hz configuration (β = −10.34, t(33) = −2.63, p = .01, 95% CI [−18.05, −2.64]). No significant effects of stimulation location were resolved for 100 Hz (β = −0.46, t(33) = −2.63, p = .91, 95% CI = [−8.61, 7.68]), bilateral 25 Hz (β = 1.59, t(17.96) = 0.45, p = .66, 95% CI = [−5.40, 8.57]), or burst stimulation (β = −0.35, t(33) = −0.09, p = .93, 95% CI = [−8.06, 7.37]).

**Figure 8.**
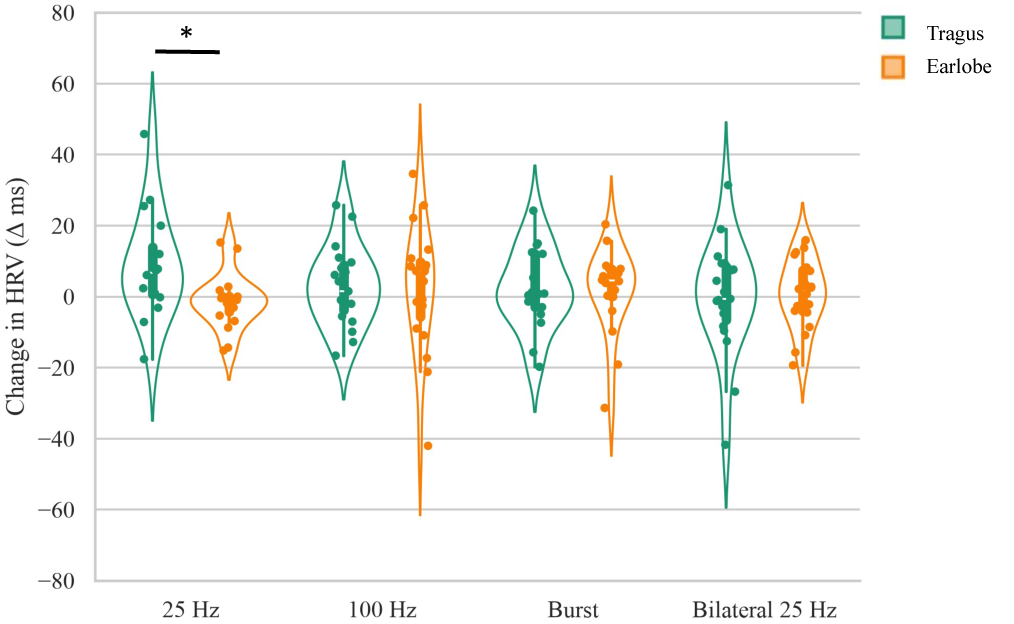
The effects of taVNS at the tragus versus the earlobe on heart rate variability for Block 1 only. Measured as change from pre-session baseline across four distinct stimulation waveforms: 25 Hz, 100 Hz, Burst, and Bilateral 25 Hz. Data are color-coded by stimulation location: Tragus (green), earlobe (orange). Individual data points represent unique participants, while the width of the violin contours illustrates the probability density of the dataset. * p<.01

### Heart Rate Variability: All Five Blocks (Post Hoc)

HRV averaged over the 60-second stimulation period across all five stimulation blocks was analyzed using linear mixed-effects models with fixed effects for stimulation location and random intercepts for subject and block within subject (Figure 9). A significant effect of stimulation location was observed only for the unilateral 25 Hz configuration (β = −1.05, t(24) =\ −2.57, p = .01, 95% CI [−1.85, −0.24]). No significant differences were detected for 100 Hz (β = −0.32, t(166.39) = 0.21, p = .84, 95% CI = [−2.71, 3.36]), bilateral 25 Hz (β = −1.26, t(170.85) = −0.86, p =, 95% CI = [−4.14, 1.61]), or burst stimulation (β = 1.68, t(168.57) = 0.95, p =0.35, 95% CI = [−1.80, 5.17]).

**Figure 9.**
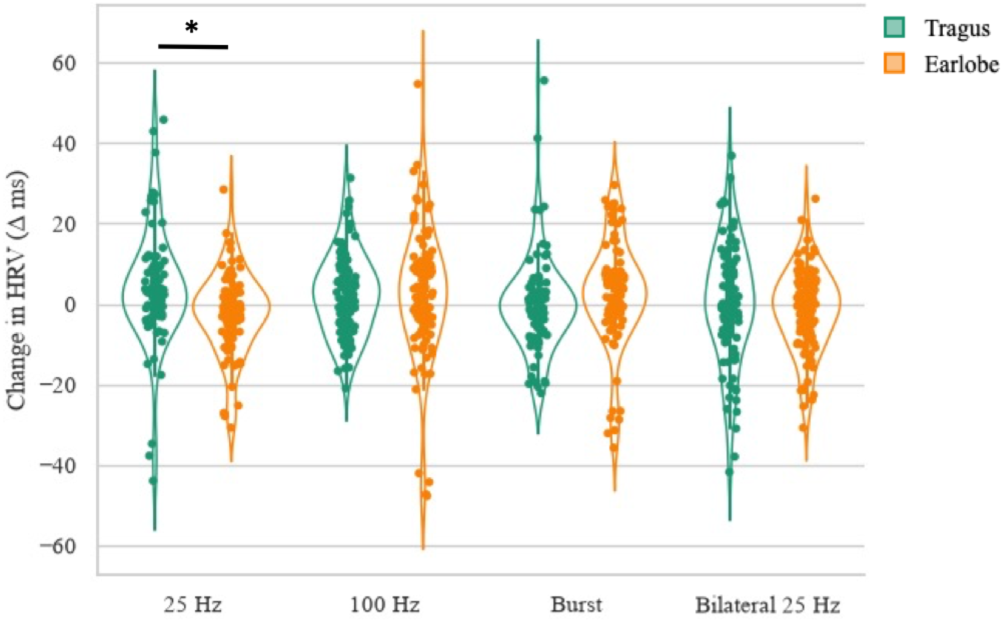
The effects of taVNS at the tragus versus the earlobe on heart rate variability for all five blocks. Measured as change from pre-session baseline across four distinct stimulation waveforms: 25 Hz, 100 Hz, Burst, and Bilateral 25 Hz. Data are color-coded by stimulation location: Tragus (green), earlobe (orange). Individual data points represent unique participants, while the width of the violin contours illustrates the probability density of the dataset. * p<.01

### Respiration: Time Averaged Effects for Block One (Primary Analysis)

Respiration was averaged over the 60-second stimulation period and analyzed using linear mixed-effects models with fixed effects for stimulation location and a random intercept for subject (Figure 10). No significant effect of stimulation location was observed for 25 Hz (β = 0.06, t(14.87) = 0.17, p = 0.86, 95% CI = [−0.63, 0.75]), 100 Hz (β = −0.56, t(39) = −1.09, p = .28, 95% CI = [−1.58, 0.45]), bilateral 25 Hz (β = 0.07, t(18.48) = −0.14, p = .63, 95% CI = [−0.67, 0.40]), or burst stimulation (β = 0.14, t(18.50) = 0.25, p = .81, 95% CI = [−0.96, 1.24]).

**Figure 10.**
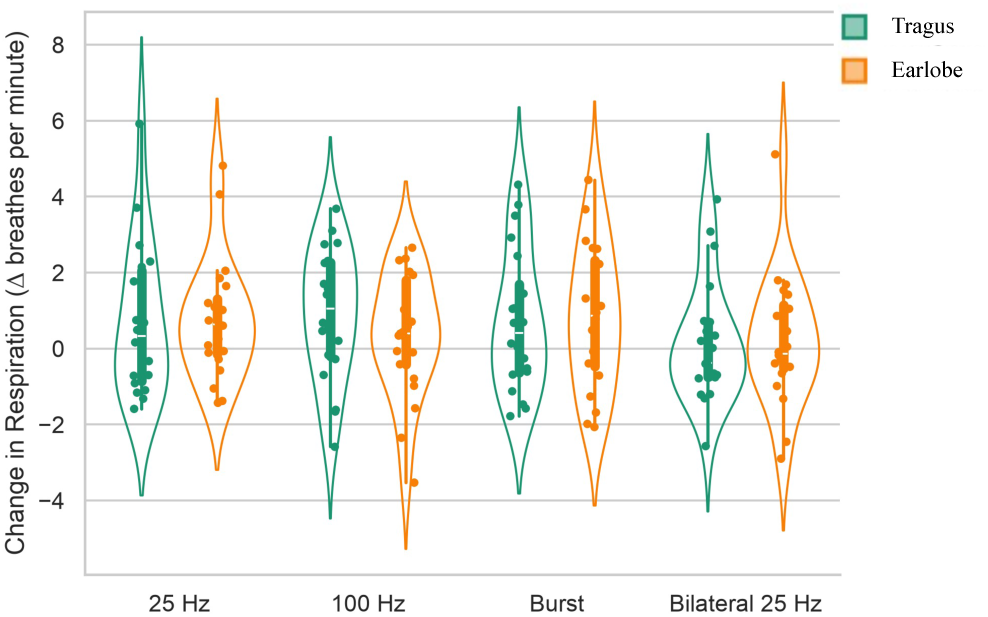
The effects of taVNS at the tragus versus the earlobe on respiration for Block 1 only. Measured as change from pre-session baseline, across four distinct stimulation waveforms: 25 Hz, 100 Hz, Burst, and Bilateral 25 Hz. Data are color-coded by stimulation location: Tragus (green) and Earlobe (orange). Individual data points represent unique participants, while the width of the violin contours illustrates the probability density of the dataset.

### Respiration: All Five Blocks (Post Hoc)

Respiration averaged over the 60-second stimulation period across all five stimulation blocks was analyzed using linear mixed-effects models with fixed effects for stimulation location and random intercepts for subject and block within subject (Figure 11). No significant effect of stimulation location was observed for any of the conditions 25 Hz (β =-0.10, t(158.12) =-0.50, p = 0.62, 95% CI = [−0.49, 0.29]), 100 Hz (β = −0.27, t(181.24) = −1.17, p = 0.24, 95% CI = [−0.72, 0.18]), bilateral 25 Hz (β = −0.13, t(181.70) = −0.73, p = 0.46, 95% CI = [−0.46, 0.21]), or burst stimulation (β = 0.11, t(170.15) = 0.52, p = 0.61, 95% CI = [−0.30, 0.51]).

**Figure 11.**
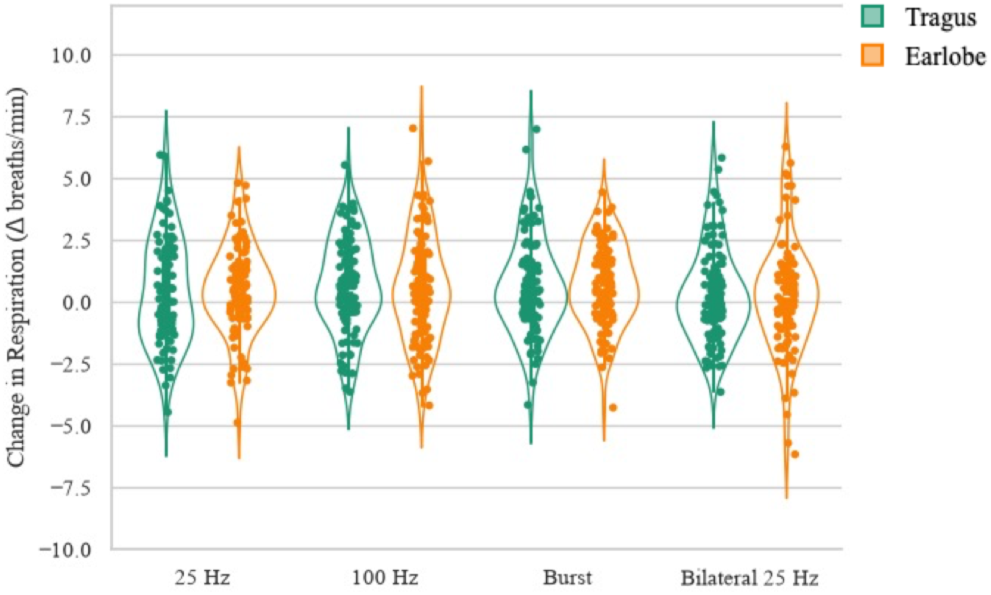
The effects of taVNS at the tragus versus the earlobe on respiration for all five blocks. Measured as change from pre-session baseline, across four distinct stimulation waveforms: 25 Hz, 100 Hz, Burst, and Bilateral 25 Hz. Data are color-coded by stimulation location: Tragus (green) and Earlobe (orange). Individual data points represent unique participants, while the width of the violin contours illustrates the probability density of the dataset.

## Discussion

This study systematically compared four taVNS stimulation configurations (left unilateral 25 Hz, 100 Hz, burst, and bilateral 25 Hz) using configuration-matched earlobe controls and a randomized within-subject design. Only unilateral left 25 Hz tragus stimulation was associated with significantly higher HRV than the matched earlobe control during both the initial stimulation block and across all five stimulation blocks, consistent with vagal target engagement. When averaged across all five blocks, unilateral left 25 Hz stimulation was also associated with higher HR relative to the matched control. Respiration did not differ between stimulation sites. Time-resolved heart rate analyses suggested location-specific differences at specific time points across the repeated blocks but not during the initial stimulation block. However, these transient effects did not alter the interpretation of the primary time-averaged analyses.

The HRV and heart rate findings reflect complementary aspects of autonomic regulation. HRV is considered to primarily reflect parasympathetic modulation, whereas mean heart rate reflects the combined influence of sympathetic and parasympathetic activity (Berntson et al., 1997; Camm et al., 1996; Shaffer and Ginsberg, 2017). Accordingly, unilateral left 25 Hz stimulation increased HRV during both the initial stimulation block and across all five blocks, consistent with an acute and sustained parasympathetic response, whereas mean heart rate differed between stimulation sites only after repeated stimulation. The delayed heart rate effect may reflect cumulative stimulation or a non-specific anticipatory response.

These findings are consistent with physiological responses to taVNS depending on stimulation configuration (Atanackov et al., 2025; Badran et al., 2019; Farmer et al., 2021; Gharabaghi and Keute, 2025; Kreisberg et al., 2021; Sclocco et al., 2020a; Sukasem et al., 2020; van Midden et al., 2024). Under the conditions tested, autonomic modulation was observed with the conventional unilateral left 25 Hz configuration, whereas significant differences were not observed for the other configurations. These findings should be interpreted carefully. Failure to detect a statistically significant difference does not establish the absence of a physiological effect, and the stimulation configurations evaluated here may produce overlapping degrees of target engagement. Moreover, autonomic responses are influenced by physiological and behavioral state. This study incorporated rigorous experimental controls - including a randomized within-subject design, configuration-matched earlobe controls, standardized cues and resting conditions, and repeated measurements; responses may differ under other behavioral contexts (Li et al., 2025) or physiological gating (Schuman-Olivier et al., 2025; Szulczewski et al., 2023). We also did not evaluate clinical efficacy. Nevertheless, establishing reproducible physiological markers of target engagement is an essential first step toward the rational optimization of taVNS.

By systematically varying waveform frequency, temporal patterning, and laterality while controlling for nonspecific sensory effects using waveform-matched earlobe stimulation, this study provides a direct test of several commonly used taVNS configurations. Taken together with prior physiological studies (Badran et al., 2018; Forte et al., 2022; Kang et al., 2024; Machetanz et al., 2021; Sclocco et al., 2020b; Taylor et al., 2025), these findings support the importance of dose-specific evaluation when characterizing the physiological effects of taVNS and selecting stimulation parameters for future mechanistic and clinical studies.

## Notes

### Competing Interest Statement

The authors have declared no competing interest.

