## Supplemental Tables 1-4 for "The Effect of Transcutaneous Auricular Vagus Nerve Stimulation (taVNS) on Cardiorespiratory Physiology: Four Waveforms Trial with Tragus vs Earlobe Stimulation"

### Supplementary Tables

| contrast | time_factor | estimate | SE | df | Lower CI | Upper CI | z ratio | p value |
|---|---|---|---|---|---|---|---|---|
| Earlobe - Tragus | stim_05 | -0.94 | 0.55 | Inf | -2.01 | 0.13 | -1.72 | 0.09 |
| Earlobe - Tragus | stim_10 | -0.67 | 0.55 | Inf | -1.74 | 0.39 | -1.24 | 0.22 |
| Earlobe - Tragus | stim_15 | -0.23 | 0.55 | Inf | -1.30 | 0.84 | -0.42 | 0.67 |
| Earlobe - Tragus | stim_20 | -1.10 | 0.55 | Inf | -2.17 | -0.04 | -2.03 | 0.04 |
| Earlobe - Tragus | stim_25 | -2.38 | 0.55 | Inf | -3.44 | -1.31 | -4.37 | <.0001 |
| Earlobe - Tragus | stim_30 | -0.76 | 0.55 | Inf | -1.82 | 0.31 | -1.39 | 0.17 |
| Earlobe - Tragus | stim_35 | -1.79 | 0.55 | Inf | -2.86 | -0.72 | -3.29 | 0.00 |
| Earlobe - Tragus | stim_40 | -0.69 | 0.55 | Inf | -1.76 | 0.38 | -1.26 | 0.21 |
| Earlobe - Tragus | stim_45 | -1.13 | 0.55 | Inf | -2.19 | -0.06 | -2.07 | 0.04 |
| Earlobe - Tragus | stim_50 | -1.35 | 0.55 | Inf | -2.42 | -0.28 | -2.48 | 0.01 |
| Earlobe - Tragus | stim_55 | -0.75 | 0.55 | Inf | -1.82 | 0.31 | -1.38 | 0.17 |
| Earlobe - Tragus | stim_60 | -1.71 | 0.55 | Inf | -2.78 | -0.64 | -3.14 | 0.00 |

Table S1. Pairwise comparisons of heart rate between tragus and earlobe stimulation across successive 5-second epochs during unilateral left 25 Hz stimulation (all five stimulation blocks).

| contrast | time_factor | estimate | SE | df | Lower CI | Upper CI | z ratio | p value |
|---|---|---|---|---|---|---|---|---|
| Earlobe - Tragus | stim_05 | -0.91 | 0.55 | Inf | -1.98 | 0.16 | -1.67 | 0.10 |
| Earlobe - Tragus | stim_10 | 0.16 | 0.55 | Inf | -0.91 | 1.23 | 0.30 | 0.77 |
| Earlobe - Tragus | stim_15 | -0.13 | 0.55 | Inf | -1.20 | 0.94 | -0.24 | 0.81 |
| Earlobe - Tragus | stim_20 | -0.20 | 0.55 | Inf | -1.26 | 0.87 | -0.36 | 0.72 |
| Earlobe - Tragus | stim_25 | -0.81 | 0.55 | Inf | -1.87 | 0.26 | -1.48 | 0.14 |
| Earlobe - Tragus | stim_30 | -0.94 | 0.55 | Inf | -2.01 | 0.13 | -1.72 | 0.09 |
| Earlobe - Tragus | stim_35 | -1.30 | 0.55 | Inf | -2.37 | -0.23 | -2.39 | 0.02 |
| Earlobe - Tragus | stim_40 | -0.63 | 0.55 | Inf | -1.70 | 0.44 | -1.16 | 0.25 |
| Earlobe - Tragus | stim_45 | -0.84 | 0.55 | Inf | -1.90 | 0.23 | -1.54 | 0.12 |
| Earlobe - Tragus | stim_50 | -0.50 | 0.55 | Inf | -1.57 | 0.57 | -0.92 | 0.36 |
| Earlobe - Tragus | stim_55 | -1.14 | 0.55 | Inf | -2.21 | -0.07 | -2.10 | 0.04 |
| Earlobe - Tragus | stim_60 | -1.36 | 0.55 | Inf | -2.42 | -0.29 | -2.49 | 0.01 |

Table S2. Pairwise comparisons of heart rate between tragus and earlobe stimulation across successive 5-second epochs during 100 Hz stimulation (all five stimulation blocks).

| contrast | time_factor | estimate | SE | df | Lower CI | Upper CI | z ratio | p value |
|---|---|---|---|---|---|---|---|---|
| Earlobe - Tragus | stim_05 | -0.43 | 0.55 | Inf | -1.50 | 0.64 | -0.79 | 0.43 |
| Earlobe - Tragus | stim_10 | -0.37 | 0.55 | Inf | -1.44 | 0.70 | -0.68 | 0.50 |
| Earlobe - Tragus | stim_15 | -0.38 | 0.55 | Inf | -1.45 | 0.69 | -0.70 | 0.48 |
| Earlobe - Tragus | stim_20 | -0.30 | 0.55 | Inf | -1.37 | 0.76 | -0.56 | 0.58 |
| Earlobe - Tragus | stim_25 | -0.02 | 0.55 | Inf | -1.09 | 1.05 | -0.04 | 0.97 |
| Earlobe - Tragus | stim_30 | -1.01 | 0.55 | Inf | -2.07 | 0.06 | -1.85 | 0.06 |
| Earlobe - Tragus | stim_35 | -0.94 | 0.55 | Inf | -2.01 | 0.13 | -1.73 | 0.08 |
| Earlobe - Tragus | stim_40 | -1.20 | 0.55 | Inf | -2.26 | -0.13 | -2.20 | 0.03 |
| Earlobe - Tragus | stim_45 | -0.69 | 0.55 | Inf | -1.76 | 0.38 | -1.27 | 0.20 |
| Earlobe - Tragus | stim_50 | -1.02 | 0.55 | Inf | -2.09 | 0.05 | -1.87 | 0.06 |
| Earlobe - Tragus | stim_55 | -0.81 | 0.55 | Inf | -1.88 | 0.26 | -1.48 | 0.14 |
| Earlobe - Tragus | stim_60 | -0.53 | 0.55 | Inf | -1.60 | 0.54 | -0.97 | 0.33 |

Table S3. Pairwise comparisons of heart rate between tragus and earlobe stimulation across successive 5-second epochs during Bilateral 25 Hz stimulation (all five stimulation blocks).

| contrast | time_factor | estimate | SE | df | Lower CI | Upper CI | z ratio | p value |
|---|---|---|---|---|---|---|---|---|
| Earlobe - Tragus | stim_05 | 0.68 | 0.55 | Inf | -0.38 | 1.75 | 1.26 | 0.21 |
| Earlobe - Tragus | stim_10 | 0.88 | 0.55 | Inf | -0.18 | 1.95 | 1.63 | 0.10 |
| Earlobe - Tragus | stim_15 | 0.69 | 0.55 | Inf | -0.38 | 1.76 | 1.27 | 0.21 |
| Earlobe - Tragus | stim_20 | 0.73 | 0.55 | Inf | -0.34 | 1.79 | 1.33 | 0.18 |
| Earlobe - Tragus | stim_25 | 0.13 | 0.55 | Inf | -0.93 | 1.20 | 0.25 | 0.81 |
| Earlobe - Tragus | stim_30 | 0.34 | 0.55 | Inf | -0.73 | 1.41 | 0.62 | 0.53 |
| Earlobe - Tragus | stim_35 | 0.04 | 0.55 | Inf | -1.02 | 1.11 | 0.08 | 0.94 |
| Earlobe - Tragus | stim_40 | 0.49 | 0.55 | Inf | -0.58 | 1.56 | 0.90 | 0.37 |
| Earlobe - Tragus | stim_45 | 0.25 | 0.55 | Inf | -0.82 | 1.32 | 0.46 | 0.65 |
| Earlobe - Tragus | stim_50 | 0.49 | 0.55 | Inf | -0.58 | 1.56 | 0.90 | 0.37 |
| Earlobe - Tragus | stim_55 | 0.28 | 0.55 | Inf | -0.79 | 1.35 | 0.52 | 0.61 |
| Earlobe - Tragus | stim_60 | -0.38 | 0.55 | Inf | -1.45 | 0.69 | -0.70 | 0.49 |

Table S4. Pairwise comparisons of heart rate between tragus and earlobe stimulation across successive 5-second epochs during burst stimulation (all five stimulation blocks).
